# A densely sampled fMRI dataset for investigating food valuation

**DOI:** 10.64898/2026.02.12.705636

**Authors:** Michiyo Sugawara, Yoko Mano, Yuhi Aoki, Koki Nakaya, Yuma Matsuda, Asako Toyama, Shinsuke Suzuki

## Abstract

Subjective valuation of food rewards guides our dietary choices and is fundamental to human health and well-being. Extensive literature in human functional magnetic resonance imaging (fMRI) studies has consistently shown that a network of reward-processing brain regions, including the ventromedial prefrontal cortex (vmPFC) and ventral striatum, encodes the subjective values of food rewards. However, the representational geometry of value signals and the mechanisms by which they are constructed in the brain remain poorly understood. This is partly because most fMRI studies on food valuation rely on small stimulus sets, yielding datasets too shallow for advanced analyses such as multi-voxel pattern analysis, and deep neural network modeling. Here, we present a densely sampled fMRI dataset wherein 31 participants provided subjective value ratings for over 500 food images across three separate days. We validate the dataset by replicating the well-established findings regarding the neural encoding of subjective value in the vmPFC and ventral striatum. We anticipate this resource will facilitate diverse studies on neural food valuation using advanced analytical methods.

## Background & Summary

Dietary choices rely on the subjective values assigned to food rewards^1,2^. Accumulating evidence in human functional magnetic resonance imaging (fMRI) studies suggests that such value signals are encoded in a network of reward-processing brain regions, including the ventromedial prefrontal cortex (vmPFC) and ventral striatum^3,4^. These regions are thought to integrate multiple types of information, including sensory pleasure, nutritional content, and health considerations, as well as various extrinsic factors^5,6^. For example, the human brain incorporates flavor^7^, oral texture^8^, nutritive attributes^9,10^, and palatability^11^, alongside price^12^ and social influence^13^ to compute the overall subjective value. Distorted neural encoding of food value signals is associated with eating disorders and obesity^14,15^. Furthermore, this value encoding has been shown to extend to other types of rewards—such as money, consumer goods, and leisure activities^16–19^—serving as a “common currency” for decision-making across different choice options. However, despite the growing literature, the representational geometry of value signals and the mechanisms by which they are constructed in the brain remain poorly understood.

Recent advances in the application of machine learning to neuroscience have opened new avenues for understanding those mechanisms. Researchers have begun to elucidate the neural computations underlying value construction by utilizing multivariate approaches such as multi-voxel pattern decoding analysis (MVPA)^20^, representational similarity analysis (RSA)^21^, and deep neural networks (DNNs)^22^ on fMRI data. MVPA has been used to decode specific content of value signals beyond simple magnitude^7,10,19^, while RSA allows for the characterization of the representational geometry of value^23^. Furthermore, deep neural networks are increasingly employed to model the non-linear transformation from raw sensory inputs, such as food images, into subjective value signals^24,25^, bridging the gap between sensory perception and decision-making.

Crucially, these advanced approaches require large amounts of data (e.g., extensive stimuli and trial repetitions) per participant to achieve stable feature representations and avoid overfitting to noise. However, most fMRI studies on food valuation use small stimulus sets; thus, conventional datasets are often too shallow to support advanced computational approaches. To address this challenge, it is necessary to collect as much data as possible within each participant, an approach known as dense within-participant sampling (or precision fMRI)^26^.

Here, we present a densely sampled fMRI dataset in which 31 participants evaluated over 500 food images across *three* separate days in an MRI scanner. Unlike previous fMRI studies that relied on repeated presentations of a limited stimulus set (e.g., 50–150 items^9–11,13,16,18,27–29^), our dataset comprises over 500 unique food images per participant. The number of food stimuli evaluated by each participant in our dataset is far greater than those in similar publicly available datasets (e.g., 108 food images for each participant^30^, and 180 food images^31^). This high-density design allows for a more robust characterization of the representational geometry of value using advanced data analyses such as MVPA decoding, RSA, and deep neural network modeling. We then technically validated the dataset by replicating the well-established finding that value signals are encoded in the vmPFC and ventral striatum (i.e., verifying that Blood-oxygenation-level-dependent signals at the time of stimulus presentation were significantly correlated with the subjective value of the food items).

## Methods

This study was approved by Hitotsubashi University IRB Committee (ID: 2025C069). All procedures were conducted following relevant ethical guidelines and regulations.

### Participants

We recruited 31 participants (11 females; age = 21.29 ± 1.87 years, mean ± SD; Body Mass Index = 20.63 ± 2.64 kg/m^2^). The sample size of 31 was determined based on our previous study^32^ in which effect sizes of the reward expectation on the vmPFC and ventral striatum activity were at minimum 0.53. With the effect size, a power analysis indicated that a sample size of at least 30 is required at a significance level of *α* = 0.05 and a statistical power of 1 - *β* = 0.80. All participants were native Japanese speakers, and pre-screened to exclude individuals with a history of neurological or psychiatric illness. Informed written consent was obtained from all participants. They received a participation fee of 9,000 Japanese Yen.

### Stimuli

We used 568 out of the 896 food images from the Food-pics dataset^33^ (Image IDs 1-568), for which nutritional information was available. These images were presented to participants in a random order.

### Experimental Task

Participants rated subjective values for 568 food images across *three* separate days (Fig. 1a). On each day, they underwent *four* fMRI runs, each consisting of 47 or 48 trials, and were asked to refrain from eating or drinking anything besides the water for 3 hours before the experiment. Note that, due to a technical issue, one participant (ID: sub-016) completed only *two* runs on day 3.

**Figure 1:**
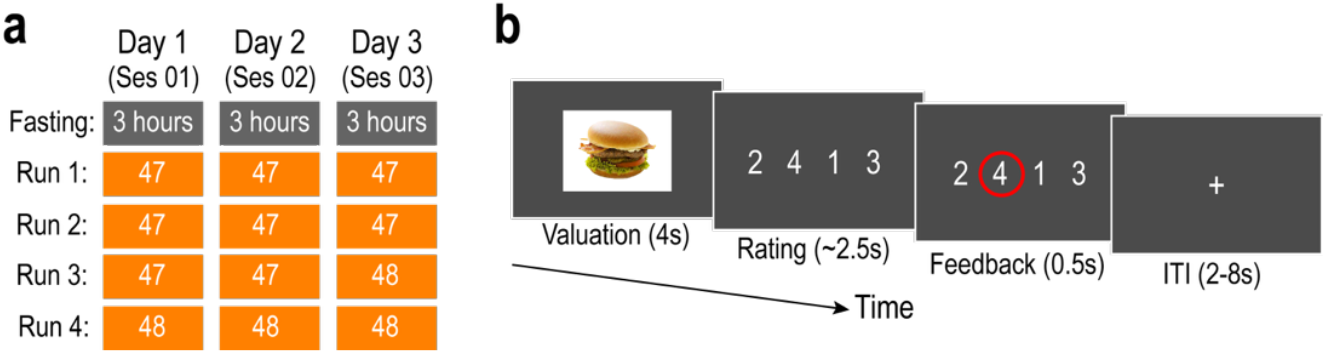
Experimental design. (a) Overall timeline. On each of the *three* days (sessions), participants completed *four* functional MRI runs following a 3-hour fast. Each run has 47 or 48 trials. Sess, session. (b) Timeline of a single trial. Participants rated the subjective value of a food item. Note that in the Rating phase, the mapping between keys and rating values was randomized across trials.

On each trial, participants rated one food image in terms of “How much do you want to eat this food?” using a 4-point Likert scale (Fig. 1b; a rating of 4 represents high value). They were first presented with a food image (Valuation phase: 4 seconds), and made a rating by pressing the corresponding key on a keypad within 2.5 seconds (Rating phase), followed by the feedback (Feedback phase: 0.5 seconds). Durations of the inter-trial-intervals (ITI) were randomly sampled from the unform distribution between 2 and 8 seconds. Here, participants were instructed to evaluate a food item during the Valuation phase, as the Rating phase was too short for decision-making.

Furthermore, to dissociate the rating value from spatial attentions and motor-related responses, the mapping between rating values (1, 2, 3, and 4) and the spatial key positions was randomized across trials.

### MRI data acquisition

Functional MRI images were collected using a 3T Siemens MAGNETOM Prisma scanner at the HIAS Brain Research Center (Hitotsubashi University, Tokyo, Japan) with a 32-channel radio frequency coil. The BOLD signal was measured using a T2*-weighted gradient-echo echo-planar imaging (EPI) sequence with the following parameters: TR = 800 ms, TE = 34.4 ms, flip angle = 52°, voxel size = 2.4 × 2.4 × 2.4 mm, matrix size = 86 × 86, 60 slices aligned to the AC–PC plane, and multiband acceleration factor = 6. For susceptibility distortion correction of the fMRI images, we also acquired a pair of spin-echo EPI “fieldmap” images with opposite phase-encoding directions (anterior-to-posterior [AP] and posterior-to-anterior [PA]) using the same geometry (FOV, matrix size, slice orientation and thickness) as the functional runs. High-resolution (0.9 mm^3^) anatomical images were acquired using a standard MPRAGE pulse sequence (TR = 2,300 ms, TE = 2.32 ms, FA = 8°).

### fMRI data preprocessing

We used *fMRIPrep* 25.2.3^34,35^, which is based on *Nipype* 1.10.0^36,37^ (RRID:SCR_002502). Following the fMRIPrep guidelines, we describe the specific details below based on the text generated by the software under a CC0 license.

#### Preprocessing of B0 inhomogeneity mappings

A total of 3 fieldmaps were found available within the input BIDS structure for this particular subject. A B0-nonuniformity map (or fieldmap) was estimated based on two (or more) echo-planar imaging (EPI) references with topup^38^ (FSL None).

#### Anatomical data preprocessing

A total of 3 T1-weighted (T1w) images were found within the input BIDS dataset. Each T1w image was corrected for intensity non-uniformity (INU) with N4BiasFieldCorrection^39^, distributed with ANTs 2.6.2^40^ (RRID:SCR_004757). The T1w-reference was then skull-stripped with a Nipype implementation of the antsBrainExtraction.sh workflow (from ANTs), using OASIS30ANTs as target template. Brain tissue segmentation of cerebrospinal fluid (CSF), white-matter (WM) and gray-matter (GM) was performed on the brain-extracted T1w using fast^41^ (FSL (version unknown), RRID:SCR_002823). An anatomical T1w-reference map was computed after registration of 3 <module ‘nipype.interfaces.image’ from ‘/app/.pixi/envs/fmriprep/lib/python3.12/site-packages/nipype/interfaces/image.py’> images (after INU-correction) using mri_robust_template^42^ (FreeSurfer 7.3.2). Brain surfaces were reconstructed using recon-all^43^ (FreeSurfer 7.3.2, RRID:SCR_001847), and the brain mask estimated previously was refined with a custom variation of the method to reconcile ANTs-derived and FreeSurfer-derived segmentations of the cortical gray-matter of Mindboggle^44^ (RRID:SCR_002438). Volume-based spatial normalization to one standard space (MNI152NLin2009cAsym) was performed through nonlinear registration with antsRegistration (ANTs 2.6.2), using brain-extracted versions of both T1w reference and the T1w template. The following templates were selected for spatial normalization and accessed with TemplateFlow^45^ (25.0.4): ICBM 152 Nonlinear Asymmetrical template version 2009c^46^ [RRID:SCR_008796; TemplateFlow ID: MNI152NLin2009cAsym].

#### Functional data preprocessing

For each of the 12 BOLD runs found per subject (across all tasks and sessions), the following preprocessing was performed. First, a reference volume was generated, using a custom methodology of *fMRIPrep*, for use in head motion correction. Head-motion parameters with respect to the BOLD reference (transformation matrices, and six corresponding rotation and translation parameters) are estimated before any spatiotemporal filtering using mcflirt^47^ (FSL). The estimated fieldmap was then aligned with rigid-registration to the target EPI (echo-planar imaging) reference run. The field coefficients were mapped on to the reference EPI using the transform. The BOLD reference was then co-registered to the T1w reference using bbregister (FreeSurfer) which implements boundary-based registration^48^. Co-registration was configured with six degrees of freedom. Several confounding time-series were calculated based on the preprocessed BOLD: framewise displacement (FD), DVARS and three region-wise global signals. FD was computed using two formulations following Power (absolute sum of relative motions^49^) and Jenkinson (relative root mean square displacement between affines^47^). FD and DVARS are calculated for each functional run, both using their implementations in Nipype (following the definitions by Power et al^49^). The three global signals are extracted within the CSF, the WM, and the whole-brain masks. Additionally, a set of physiological regressors were extracted to allow for component-based noise correction^50^ (CompCor). Principal components are estimated after high-pass filtering the preprocessed BOLD time-series (using a discrete cosine filter with 128s cut-off) for the two CompCor variants: temporal (tCompCor) and anatomical (aCompCor). tCompCor components are then calculated from the top 2% variable voxels within the brain mask. For aCompCor, three probabilistic masks (CSF, WM and combined CSF+WM) are generated in anatomical space. The implementation differs from that of Behzadi et al. in that instead of eroding the masks by 2 pixels on BOLD space, a mask of pixels that likely contain a volume fraction of GM is subtracted from the aCompCor masks. This mask is obtained by dilating a GM mask extracted from the FreeSurfer’s aseg segmentation, and it ensures components are not extracted from voxels containing a minimal fraction of GM. Finally, these masks are resampled into BOLD space and binarized by thresholding at 0.99 (as in the original implementation). Components are also calculated separately within the WM and CSF masks. For each CompCor decomposition, the k components with the largest singular values are retained, such that the retained components’ time series are sufficient to explain 50 percent of variance across the nuisance mask (CSF, WM, combined, or temporal). The remaining components are dropped from consideration.

The head-motion estimates calculated in the correction step were also placed within the corresponding confounds file. The confound time series derived from head motion estimates and global signals were expanded with the inclusion of temporal derivatives and quadratic terms for each^51^. Frames that exceeded a threshold of 0.5 mm FD or 1.5 standardized DVARS were annotated as motion outliers. Additional nuisance timeseries are calculated by means of principal components analysis of the signal found within a thin band (crown) of voxels around the edge of the brain, as proposed by (Patriat, Reynolds, and Birn 2017)^52^. All resamplings can be performed with a single interpolation step by composing all the pertinent transformations (i.e. head-motion transform matrices, susceptibility distortion correction when available, and co-registrations to anatomical and output spaces). Gridded (volumetric) resamplings were performed using nitransforms, configured with cubic B-spline interpolation.

### Data Records

All data are organized according to the Brain Imaging Data Structure (BIDS) ^53^ and are available on the OpenNeuro database (https://openneuro.org/datasets/ds007267)^54^. Each participant is assigned a unique identifier (e.g., sub-001), and demographic information is provided in the *participants*.*tsv* file. The dataset consists of *three* sessions per participant, corresponding to the *three* experimental days (labeled ses-01, ses-02, and ses-03). Each session directory contains *three* subdirectories: *anat, fmap*, and *func*, housing anatomical, field-map, and functional images, respectively. Typically, the *func* directory contains four experimental runs; however, due to a technical issue, one participant (sub-016) completed only *two* runs in the *third* session (ses-03).

Detailed behavioral data are provided in event files (e.g., *sub-001_ses-01_task-food_run-01_events*.*tsv*) located within the *func* directory. These event files contain trial-wise information, including stimulus properties (the food image presented, along with its visual and nutritional features); participant responses (the subjective ratings submitted by the participant); and timing (onsets and durations for the Valuation, Rating, and Feedback phases, as well as the timing of key-press responses).

### Technical Validation

#### Basic checks

We first examined the proportion of miss trials in which participants failed to make a rating within the time limit. The error rates were low across the cohort, with a median of 0.013 (25th percentile = 0.002; and 75th percentile = 0.060). The rate was less than 0.10 in 28 out of 31 participants, and remained below 0.15 for all participants, indicating that subjects maintained high levels of attention throughout the experimental task.

To assess the quality of the fMRI data, we evaluated the temporal signal-to-noise ratio (tSNR) and framewise displacement (FD). First, to quantify signal strength, we calculated the tSNR (defined as the temporal mean divided by the temporal standard deviation per voxel) of the preprocessed data. We visualized the average tSNR across participants on the cortical surface (Fig. 2a, left) and specifically examined regions of interest (ROIs) implicated in encoding food value signals^53–57^: the vmPFC and ventral striatum. These ROIs were defined as 8-mm spheres centered on peak coordinates extracted from the previous meta-analysis^4^ ([x, y, z] = [-2, 40, -6] for vmPFC; [10, 12, -10] for left ventral striatum; and [-10, 8, -6] for right ventral striatum) (Fig. 2b). Furthermore, to assess volume-to-volume head movement, we calculated the average FD value for each participant^49,55^ (Fig. 2a, right). The median FD across participants was 0.104 (25th percentile = 0.083; 75th percentile = 0.153). This distribution was comparable to that observed in other datasets^56–60^, indicating that head motion remained within an acceptable range.

**Figure 2:**
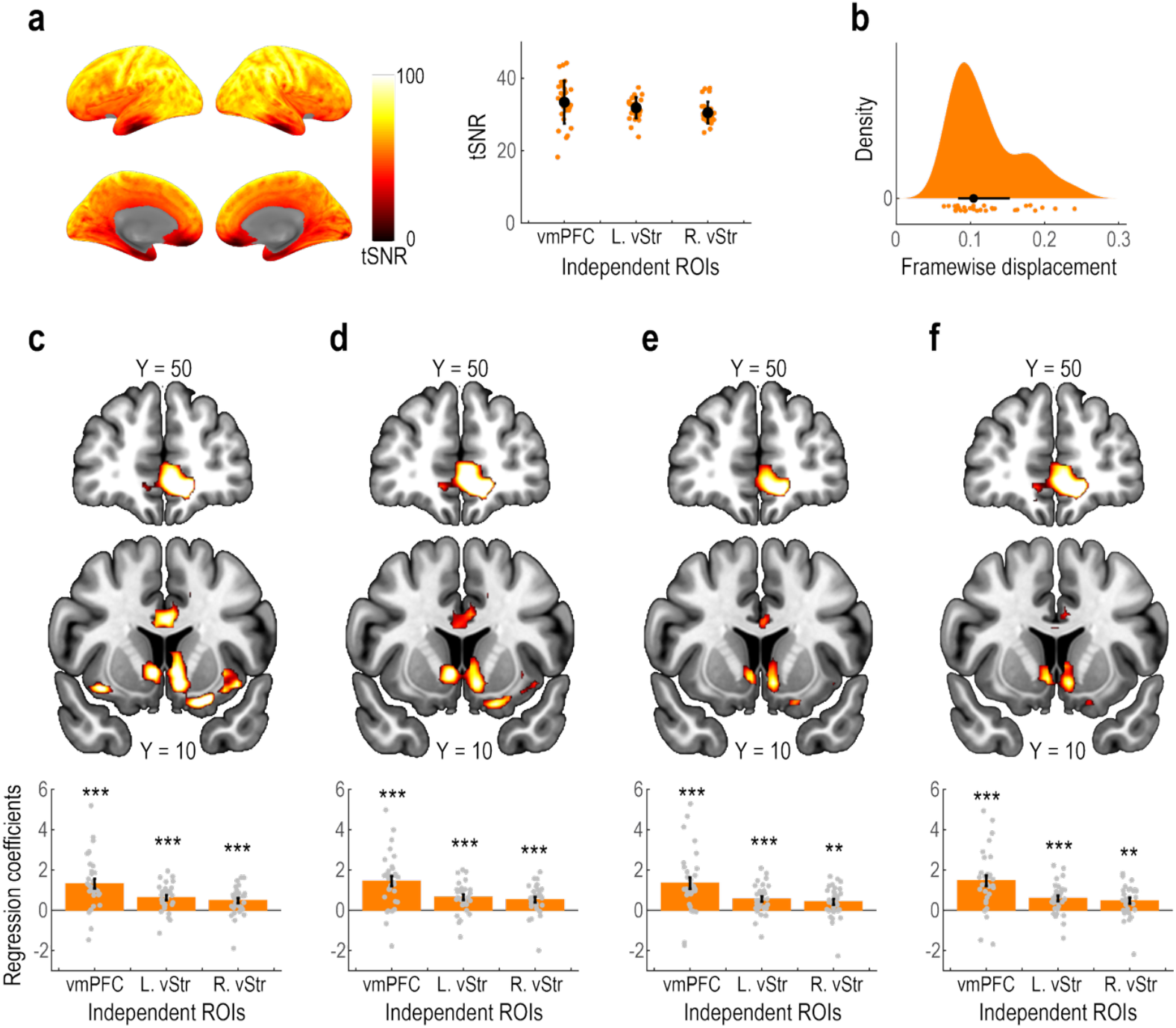
fMRI data analyses for the technical validation. (a) Temporal signal to noise ratio (tSNR). *Left*: cortical surface map showing the mean tSNR value across participants in each voxel. *Right*: tSNR values in independently defined regions of interest. Each point denotes one participant. Black points and vertical bars indicate the mean and SD across participants. vmPFC, ventromedial prefrontal cortex; L.vStr, left ventral striatum; and R.vStr, right ventral striatum. (b) Framewise displacement (FD). The density plot shows the distribution of mean FD values across participants. Each point denotes one participant. The black point and horizontal bar indicate the median and 25%-75% quantiles across participants. (c) Neural correlates of subjective value (GLM1). *Top* and *middle*: whole-brain statistical parametric map showing BOLD signal parametrically modulated by the rating value during the Valuation phase. The activation map is thresholded at *p* < 0.001 (uncorrected) for display purposes. *Bottom*: Independent ROI analysis. Bar plots show the regression coefficients of subjective value (Mean ± SEM across participants) in independently defined regions of interest. Each gray point depicts one participant. \*\*\**p* < 0.001 and \*\**p* < 0.01. (d) Neural correlates of subjective value after controlling for color intensities of the food images (GLM2). The format is the same as (c). (e) Neural correlates of subjective value after controlling for nutrient contents of the food images (GLM3). The format is the same as (c). (f) Neural correlates of subjective value after controlling for both color intensities and nutrient contents (GLM4). The format is the same as (c).

#### Replication of previous findings

We replicated well-established findings that value signals for food rewards are encoded in a network of reward-processing regions, including the vmPFC and ventral striatum^49,58,59^. To this end, we performed conventional General Linear Model (GLM) analyses on the smoothed (6-mm FWHM Gaussian kernel) and high-pass filtered (128 s cutoff) fMRI data. GLM1 included the following regressors for each participant: *three* boxcar functions for the Valuation (duration = 4 s), Response (duration = reaction time), and Feedback phases (duration = 0.5 s); a regressor for missed trials; and a stick function for the timing of the key press. The Valuation regressor included a parametric modulator encoding the subjective value (i.e., rating value) of the presented food image. All regressors were convolved with a canonical hemodynamic response function (HRF). Additionally, six motion-correction parameters and indicator regressors for volumes exhibiting excessive framewise displacement (FD > 0.9 mm) were included to account for head movement artifacts^49,55,61^. The contrast map for subjective value was estimated for each participant and submitted to a random-effect group-level analysis. We also tested three GLMs including additional parametric modulators during the Valuation phase: color intensities (red, green, blue) in GLM2; nutrient content (protein, fat, carbohydrate) in GLM3; and both color and nutrients in GLM4.

Consistent with prior studies^3,4^, the whole-brain analysis revealed that the vmPFC and ventral striatum significantly encoded value signals (Fig. 2c; *p* < 0.05 corrected for multiple comparisons at the cluster level; cluster-forming threshold *p* < 0.001). We further confirmed this using independent ROI analyses. BOLD signals in the vmPFC and ventral striatum were significantly correlated with subjective value at the time of valuation (*t*(30) = 5.47, *p* < 0.001 for vmPFC; *t*(30) = 5.15, *p* < 0.001 for left ventral striatum; *t*(30) = 3.80, *p* < 0.001 for right ventral striatum). Importantly, food value encoding remained significant even after controlling for potential confounds (Fig. 2def). ROI analyses yielded significant results after controlling for color intensity (all *ps* < 0.001), nutrient content (all *ps* < 0.01), and both factors simultaneously (all *ps* < 0.01). These results collectively validate our experimental design by robustly replicating established neural value signatures.

## Acknowledgements

We thank Kimiko Watanabe for her assistance with participant recruitment.

## Data availability

All data are organized according to the Brain Imaging Data Structure (BIDS) ^53^ and are available on the OpenNeuro database (https://openneuro.org/datasets/ds007267)^54^.

## Code availability

The code for data analysis is available at the following URL: https://github.com/szkshnsk/A-densely-sampled-fMRI-dataset-for-investigating-food-valuation

## References

1. Rangel, A., Camerer, C. & Montague, P. R. A framework for studying the neurobiology of value-based decision making. Nat. Rev. Neurosci. 9, 545–556 (2008).

2. Ruff, C. C. & Fehr, E. The neurobiology of rewards and values in social decision making. Nat. Rev. Neurosci. 15, 549–562 (2014).

3. Bartra, O., McGuire, J. T. & Kable, J. W. The valuation system: a coordinate-based meta-analysis of BOLD fMRI experiments examining neural correlates of subjective value. Neuroimage 76, 412–427 (2013).

4. Clithero, J. A. & Rangel, A. Informatic parcellation of the network involved in the computation of subjective value. Soc. Cogn. Affect. Neurosci. 9, 1289–1302 (2014).

5. Motoki, K. & Suzuki, S. Extrinsic factors underlying food valuation in the human brain. Front. Behav. Neurosci. 14, 131 (2020).

6. Suzuki, S. Constructing value signals for food rewards: determinants and the integration. Current Opinion in Behavioral Sciences 46, 101178 (2022).

7. Howard, J. D., Gottfried, J. A., Tobler, P. N. & Kahnt, T. Identity-specific coding of future rewards in the human orbitofrontal cortex. Proc. Natl. Acad. Sci. U. S. A. 112, 5195–5200 (2015).

8. Khorisantono, P. A. et al. A neural mechanism in the human orbitofrontal cortex for preferring high-fat foods based on oral texture. J. Neurosci. 43, 8000–8017 (2023).

9. DiFeliceantonio, A. G. et al. Supra-additive effects of combining fat and carbohydrate on food reward. Cell Metab. 28, 33–44.e3 (2018).

10. Suzuki, S., Cross, L. & O’Doherty, J. P. Elucidating the underlying components of food valuation in the human orbitofrontal cortex. Nat. Neurosci. 20, 1780–1786 (2017).

11. Plassmann, H., O’Doherty, J. P. & Rangel, A. Appetitive and aversive goal values are encoded in the medial orbitofrontal cortex at the time of decision making. J. Neurosci. 30, 10799–10808 (2010).

12. Plassmann, H., O’Doherty, J., Shiv, B. & Rangel, A. Marketing actions can modulate neural representations of experienced pleasantness. Proceedings of the National Academy of Sciences 105, 1050–1054 (2008).

13. Nook, E. C. & Zaki, J. Social norms shift behavioral and neural responses to foods. J. Cogn. Neurosci. 27, 1412–1426 (2015).

14. Foerde, K., Steinglass, J. E., Shohamy, D. & Walsh, B. T. Neural mechanisms supporting maladaptive food choices in anorexia nervosa. Nat. Neurosci. 18, 1571–1573 (2015).

15. Seabrook, L. T. & Borgland, S. L. The orbitofrontal cortex, food intake and obesity. J. Psychiatry Neurosci. 45, 304–312 (2020).

16. Chib, V. S., Rangel, A., Shimojo, S. & O’Doherty, J. P. Evidence for a common representation of decision values for dissimilar goods in human ventromedial prefrontal cortex. J. Neurosci. 29, 12315–12320 (2009).

17. Gross, J. et al. Value Signals in the Prefrontal Cortex Predict Individual Preferences across Reward Categories. J. Neurosci. 34, 7580–7586 (2014).

18. Levy, D. J. & Glimcher, P. W. Comparing apples and oranges: using reward-specific and reward-general subjective value representation in the brain. J. Neurosci. 31, 14693–14707 (2011).

19. McNamee, D., Rangel, A. & O’Doherty, J. P. Category-dependent and category-independent goal-value codes in human ventromedial prefrontal cortex. Nat. Neurosci. 16, 479–485 (2013).

20. Haynes, J.-D. A primer on pattern-based approaches to fMRI: Principles, pitfalls, and perspectives. Neuron 87, 257–270 (2015).

21. Kriegeskorte, N., Mur, M. & Bandettini, P. Representational similarity analysis - connecting the branches of systems neuroscience. Front. Syst. Neurosci. 2, 4 (2008).

22. Pham, T. Q., Matsui, T. & Chikazoe, J. Evaluation of the hierarchical correspondence between the human brain and artificial neural networks: A review. Biology (Basel) 12, 1330 (2023).

23. Avery, J., Carrington, M., Liu, A. & Martin, A. Representation of naturalistic food categories in the human brain. J. Vis. 22, 3426 (2022).

24. Iigaya, K. et al. Neural mechanisms underlying the hierarchical construction of perceived aesthetic value. Nat. Commun. 14, 1–19 (2023).

25. Pham, T. Q. et al. Vision-to-value transformations in artificial neural networks and human brain. bioRxiv (2021) doi:10.1101/2021.03.18.435929.

26. Gratton, C. & Braga, R. M. Dense phenotyping of human brain network organization using precision fMRI. Annu. Rev. Psychol. (2025) doi:10.1146/annurev-psych-032825-032920.

27. Takehana, A. et al. Healthy dietary choices involve prefrontal mechanisms associated with long-term reward maximization but not working memory. Cereb. Cortex 34, (2024).

28. Enax, L., Hu, Y., Trautner, P. & Weber, B. Nutrition labels influence value computation of food products in the ventromedial prefrontal cortex: Impact of Nutrional Labels on Valuation in vmPFC. Obesity (Silver Spring) 23, 786–792 (2015).

29. Hare, T. A., Camerer, C. F. & Rangel, A. Self-control in decision-making involves modulation of the vmPFC valuation system. Science 324, 646–648 (2009).

30. Tomova, L. et al. MRI data of 40 adult participants in response to a cue induced craving task following food fasting, social isolation and baseline (within-subject design). MRI data of 40 adult participants in response to a cue induced craving task following food fasting, social isolation and baseline (within-subject design) Openneuro 10.18112/OPENNEURO.DS003242.V1.0.0 (2020).

31. Courtney, A., PeConga, E., Wagner, D. & Rapuano, K. Calorie-labeled food cues. Calorie-labeled food cues Openneuro 10.18112/OPENNEURO.DS001534.V1.1.0 (2019).

32. Ohnishi, K., Sugawara, M., Mano, Y. & Suzuki, S. Neural mechanisms underlying reward processing and social cognition: A replication study with a Japanese sample. PLoS One 20, e0328424 (2025).

33. Blechert, J., Lender, A., Polk, S., Busch, N. A. & Ohla, K. Food-pics_extended-an image database for experimental research on eating and appetite: Additional images, normative ratings and an updated review. Front. Psychol. 10, 307 (2019).

34. Esteban, O. et al. fMRIPrep: a robust preprocessing pipeline for functional MRI. Nat. Methods 16, 111–116 (2019).

35. Markiewicz, C. J., Esteban, O., Goncalves, M., Poldrack, R. A. & Gorgolewski, K. J. FMRIPrep: A Robust Preprocessing Pipeline for Functional MRI. (Zenodo, 2025). doi:10.5281/ZENODO.852659.

36. Gorgolewski, K. et al. Nipype: a flexible, lightweight and extensible neuroimaging data processing framework in python. Front. Neuroinform. 5, 13 (2011).

37. Esteban, O. et al. Nipy/Nipype: 1.9.1. (Zenodo, 2025). doi:10.5281/ZENODO.596855.

38. Andersson, J. L. R., Skare, S. & Ashburner, J. How to correct susceptibility distortions in spin-echo echo-planar images: application to diffusion tensor imaging. Neuroimage 20, 870–888 (2003).

39. Tustison, N. J. et al. N4ITK: improved N3 bias correction. IEEE Trans. Med. Imaging 29, 1310–1320 (2010).

40. Avants, B. B., Epstein, C. L., Grossman, M. & Gee, J. C. Symmetric diffeomorphic image registration with cross-correlation: evaluating automated labeling of elderly and neurodegenerative brain. Med. Image Anal. 12, 26–41 (2008).

41. Zhang, Y., Brady, M. & Smith, S. Segmentation of brain MR images through a hidden Markov random field model and the expectation-maximization algorithm. IEEE Trans. Med. Imaging 20, 45–57 (2001).

42. Reuter, M., Rosas, H. D. & Fischl, B. Highly accurate inverse consistent registration: a robust approach. Neuroimage 53, 1181–1196 (2010).

43. Dale, A. M., Fischl, B. & Sereno, M. I. Cortical surface-based analysis. I. Segmentation and surface reconstruction. Neuroimage 9, 179–194 (1999).

44. Klein, A. et al. Mindboggling morphometry of human brains. PLoS Comput. Biol. 13, e1005350 (2017).

45. Ciric, R. et al. TemplateFlow: FAIR-sharing of multi-scale, multi-species brain models. Nat. Methods 19, 1568–1571 (2022).

46. Fonov, V. S., Evans, A. C., McKinstry, R. C., Almli, C. R. & Collins, D. L. Unbiased nonlinear average age-appropriate brain templates from birth to adulthood. Neuroimage 47, S102 (2009).

47. Jenkinson, M., Bannister, P., Brady, M. & Smith, S. Improved optimization for the robust and accurate linear registration and motion correction of brain images. Neuroimage 17, 825–841 (2002).

48. Greve, D. N. & Fischl, B. Accurate and robust brain image alignment using boundary-based registration. Neuroimage 48, 63–72 (2009).

49. Power, J. D. et al. Methods to detect, characterize, and remove motion artifact in resting state fMRI. Neuroimage 84, 320–341 (2014).

50. Behzadi, Y., Restom, K., Liau, J. & Liu, T. T. A component based noise correction method (CompCor) for BOLD and perfusion based fMRI. Neuroimage 37, 90–101 (2007).

51. Satterthwaite, T. D. et al. An improved framework for confound regression and filtering for control of motion artifact in the preprocessing of resting-state functional connectivity data. Neuroimage 64, 240–256 (2013).

52. Patriat, R., Reynolds, R. C. & Birn, R. M. An improved model of motion-related signal changes in fMRI. Neuroimage 144, 74–82 (2017).

53. Gorgolewski, K. J. et al. The brain imaging data structure, a format for organizing and describing outputs of neuroimaging experiments. Sci. Data 3, 160044 (2016).

54. Sugawara, M. et al. A densely sampled fMRI dataset for investigating food valuation. Openneuro 10.18112/OPENNEURO.DS007267.V1.0.0 (2026).

55. Power, J. D., Schlaggar, B. L. & Petersen, S. E. Recent progress and outstanding issues in motion correction in resting state fMRI. Neuroimage 105, 536–551 (2015).

56. Wang, L.-S. et al. An fMRI hyperscanning dataset on cooperation and competition during strategic interactions. Sci. Data 12, 1414 (2025).

57. Wang, B., Zhang, X., Zhang, L. & Kong, X.-Z. A naturalistic fMRI dataset in response to public speaking. Sci. Data 12, 659 (2025).

58. Wang, S. et al. An fMRI dataset for concept representation with semantic feature annotations. Sci. Data 9, 721 (2022).

59. Jung, H. et al. Spacetop: A multimodal fMRI dataset unifying naturalistic processes with a rich array of experimental tasks. Sci. Data 12, 1465 (2025).

60. Guo, T., Liu, X., Chen, M., Fu, Y. & Guo, T. An fMRI dataset for investigating language control and cognitive control in bilinguals. Sci. Data 12, 889 (2025).

61. Siegel, J. S. et al. Statistical improvements in functional magnetic resonance imaging analyses produced by censoring high-motion data points: Censoring High Motion Data in fMRI. Hum. Brain Mapp. 35, 1981–1996 (2014).

